# LAMELLAR THICKNESS MEASUREMENTS IN NORMAL AND OSTEOGENESIS IMPERFECTA HUMAN BONE, with development of a method of automated thickness averaging to simplify quantitation

**DOI:** 10.1101/2022.05.30.493917

**Authors:** J Chow, N Ryan, SJ Shefelbine, F Shapiro

## Abstract

**Purpose:** Lamellar bone that forms in moderate and severe osteogenesis imperfecta (OI) is often composed of structurally irregular lamellae compared to those in normal bone. Polarization light microscopy (PLM) demonstrates lamellar bone well but has rarely been used for quantitative studies; information available on normal bone lamellae tends to be variable and studies specifically assessing OI bone lamellae have not been done. We report on PLM histomorphometry quantifying bright and dark lamellar thicknesses in normal and OI bone. Manual measurements of individual lamellar thicknesses have been made on histologic sections using the cellSens image analysis system; in an effort to augment the number of measurements we also developed a method of automated thickness averaging in quantifying regions of lamellae.

**Methods:** Femoral and tibial cortical bone fragments from 5 individuals 5 – 26 years old (without molecular bone disorders) and 8 individuals 5 – 16 years old with progressively deforming (Sillence III) OI were obtained. The fragments were decalcified, infiltrated in JB4 solution, embedded in JB4 plastic, sectioned at 5μ thickness and stained with 1% toluidine blue for light and polarizing microscopy. *Manual measurements:* Strict criteria for measurement, primarily to eliminate oblique lamellae, included accumulations of 16-20 bright and dark lamellae under PLM with a relatively narrow range of thicknesses, flattened elliptical osteocytes along the longitudinal axis of the lamellae and canaliculi passing from the walls of the osteocyte lacunae at right angles to the lamellae. Histomorphometric measurements of bright and dark lamellae by PLM were made at 20X magnification. *Automated measurements:* A script for automated measurement of average lamellar thicknesses from PLM images was developed in MATLAB (Mathworks, Natick, MA) to make measurement faster and less subjective. The script isolates a region from an image for measurement and marks each pixel as either bright or dark based on a local average intensity threshold. It then takes multiple pixel measurements along the length of the lamellae in the image and returns the average thickness of each in μm.

**Results:** *1. OI bone mean lamellar thickness values are always less than those in normal bone*. The mean value for all OI bright and dark lamellae combined is 1.80 ± 0.72 μm and the value in normal bone is 2.54 ± 0.92 μm. *2. Mean value for the bright lamellae is less than that for the dark lamellae in both normal and OI bone*. The mean value for bright lamellae in OI is 1.47 ± 0.53 μm and for dark lamellae 2.18 ± 0.72 μm; in normal bone the mean value for bright lamellae is 2.06 ± 0.54 μm and for dark lamellae 3.07 ± 0.96 μm. The differences are statistically significant: between groups of normal and OI lamellae (p<0.001), normal and OI light bands (p<0.001), and normal and OI dark bands (p<0.001). *3. Ratio of mean values for bright/dark lamellar thicknesses is the same in OI and normal bone*. The ratio in OI bone is 0.67 (range: 0.54 – 0.83) and in normal bone 0.67 (range: 0.60 – 0.88). *4. Validation of automated vs. manual datasets:* For each lamella in the validation dataset, the percent difference between the automated and manual measurements was calculated. The mean of the absolute values of these percent differences was 18.9%, a statistically non-significant difference (p = 0.0518).

**Discussion and conclusions:** Lamellar bone that forms in moderate and severe OI is composed of thinner and less regular lamellae than those in normal bone. i) PLM histomorphometry shows mean lamellar thicknesses (bright and dark merged) are statistically significantly decreased in OI compared to normal bone as are bright and dark lamellar thicknesses measured independently. ii) The automated method can be adapted readily to the assessment process for lamellar thicknesses and is, most likely, more accurate since it averages a greatly increased number of measurements per individual lamella. iii) Lamellar thickness measurements can be helpful in assessing the effect of specific collagen mutations on OI bone synthesis and warrant inclusion in both research and clinical histomorphometric assessments.

## INTRODUCTION

Collagen matrices are aligned preponderantly in parallel, or more accurately, unidirectional fashion in normally developed cortical bone. The basic structural components in bone were recognized over 300 years ago by Havers (1691) who identified what he referred to as laminae along the longitudinal axis in bone and van Leeuwenhoek (1693) who first observed osteons. The structure of bone was more clearly defined in the mid 1800s as light microscopy techniques were developed; observers including Kölliker (1850) and Tomes and de Morgan (1853) defined the development of bone as a tissue recognizing the circular layers of lamellae around central longitudinal intra-cortical blood vessels forming osteons and now recognized as the main structural element of cortical bone. (Todd and Bowman in 1845 named the osteonal lamellar layers surrounding a central vessel as the Haversian system.) Polarization microscopy was discovered in 1828 by William Nicol (see Cheshire, 1923) and in time was used to assess bone. Von Ebner (1875) used polarization to highlight the lamellar structure of bone showing alternating bright and dark layers that he attributed to alternating change in the orientation of the fibrillar bundles. Ziegler (1906) also used polarization but attributed the alternating layers to differences in their components. This layering of the lamellae has thus been of scientific interest and investigation for over 150 years. Gebhardt (1906) wrote in detail on the lamellae and their three-dimensional organization as revealed by polarization microscopy. Representation of the long axes of adjacent lamellae as being positioned at right angles to one another, in what is referred to as an orthogonal array, has been attributed to Gebhardt. Reference to his original 1906 paper, however, shows a much more detailed and sophisticated description; the basic orthogonal pattern was included (although that term was not used) but the alternating layers were demonstrated to align as adjacent layers of fibrils rotating in opposite directions to one another although only occasionally angling to such an extent that they were at right angles. Over the past several years it has been recognized that the lamellae of cortical bone osteons, when assessed in three-dimensional fashion and at higher levels of structural resolution, are organized into a twisted plywood-like arrangement rather than strictly parallel sets of lamellae running rigidly along the longitudinal and transverse axes (Giraud-Guille, 1988). Wagermaier et al (2006) recognized that not only are individual collagen fibrils twisted as a helix; there is a helicoidal spiral twisting arrangement of collections of fibrils in osteons as well with varying degrees of angulation. It is now recognized that the fibrillar (collagenous) organization of bone shows both orthogonal and helicoidal architecture (Neville, 1993). Buss et al (2022) have observed that at 9 of the currently recognized 12 hierarchical layers of bone structure, the components are organized into the spiraling / twisting / coiled conformation.

There have been varying interpretations of the alternating bright and dark lamellae based on study at higher levels of structural resolution than PLM, in particular transmission and scanning electron microscopy (Marotti, 1993; Reznikov et al, 2014a; Mitchell and van Heteren, 2016). PLM, even though available for such a long period of time, has been used to assess bone primarily in a qualitative fashion to differentiate woven from lamellar bone but there has been very little quantitative work done in normal bone and essentially none regarding the lamellar structure in OI bone. Cortical bone development in OI patients, while imperfect due to the underlying collagen mutations, still progresses to a lamellar arrangement in progressively deforming and less affected autosomal dominant variants. This progression occurs once a sufficient amount of woven bone has been synthesized by what we refer to as mesenchymal osteoblasts to serve as a scaffold for surface osteoblasts that align in single array on the woven scaffold and synthesize lamellar bone (Shapiro, 2008; Marotti, 2010; Shapiro and Wu, 2019). The lamellar segments in OI, however, are often structured into a mosaic collection of short segments oriented in random directions to one another. While studying OI bone at multiple levels of structural resolution we also noted qualitatively that the individual lamellar thicknesses often appeared different from those in normal bone (Shapiro et al, 2021) often showing thinner layers with structural irregularity. Lamellar thicknesses in OI bone have not been documented in any controlled fashion and, theoretically, could have biomechanical implications. In this study we quantify lamellar thicknesses in normal human and OI bone. While deeper understanding of the three-dimensional organization of the lamellae and the specific constituents of their bright and dark layers will come from studies at higher levels of resolution, we demonstrate that quantitative PLM (via histomorphometric techniques) can enhance other correlative studies and, by itself, can provide valuable information for basic research and clinical application. In this study manual measurements were made of the individual lamellar thicknesses in normal and OI bone following which an automated algorithm has been developed to streamline these measurements for wider application.

## MATERIALS AND METHODS

### a) Bone tissues studied

*i) Normal bone* (*bone without underlying molecular abnormalities)*. Bone specimens without underlying molecular abnormalities (referred to as normal bone) were obtained from 5 individuals (5 – 26 years of age) undergoing orthopedic surgical corrective procedures. The tissue studied was femoral or tibial cortical diaphyseal bone removed while obtaining correction of deformity that would otherwise have been discarded. The bone tissue was either fully normal (representing tissue removed from the longer limb during bone length equalization procedures) or non-pathologic (representing bone from patients with cerebral palsy or developmental dysplasia of the hip undergoing deformity correction by osteotomy). The sources of the normal bone tissue, labeled CS1 to 5, are listed in table 1. *ii) Osteogenesis imperfecta bone*. Bone specimens from 8 patients (5 – 16 years of age) with OI were obtained from intact lower extremity long bones (femur or tibia) at time of osteotomy and intramedullary rod insertion to correct deformity. Osteotomies were done electively on intact but deformed bones; none of the bone tissue was obtained at time of or shortly after acute fractures and tissue removed as part of correction otherwise would have been discarded. The OI patients were categorized using the Sillence terminology (Marini et al, 2017). Bone was obtained from progressively deforming Sillence III OI patients with femoral or tibial deformities severe enough to warrant surgical intervention by osteotomy (to straighten deformity) followed by intramedullary rod insertion. Neither Sillence II (lethal perinatal) or Sillence I (benign dominant) patients were included; the Sillence II group forms only woven bone with no significant accumulations of lamellar bone and Sillence I patients, where bone is lamellar in structure, are relatively mildly affected and rarely require surgical intervention. The sources of OI bone tissue, labeled CS 6 to 13, are listed in table 1. The investigation was approved by the hospital Institutional Review Boards.

**Table 1.**
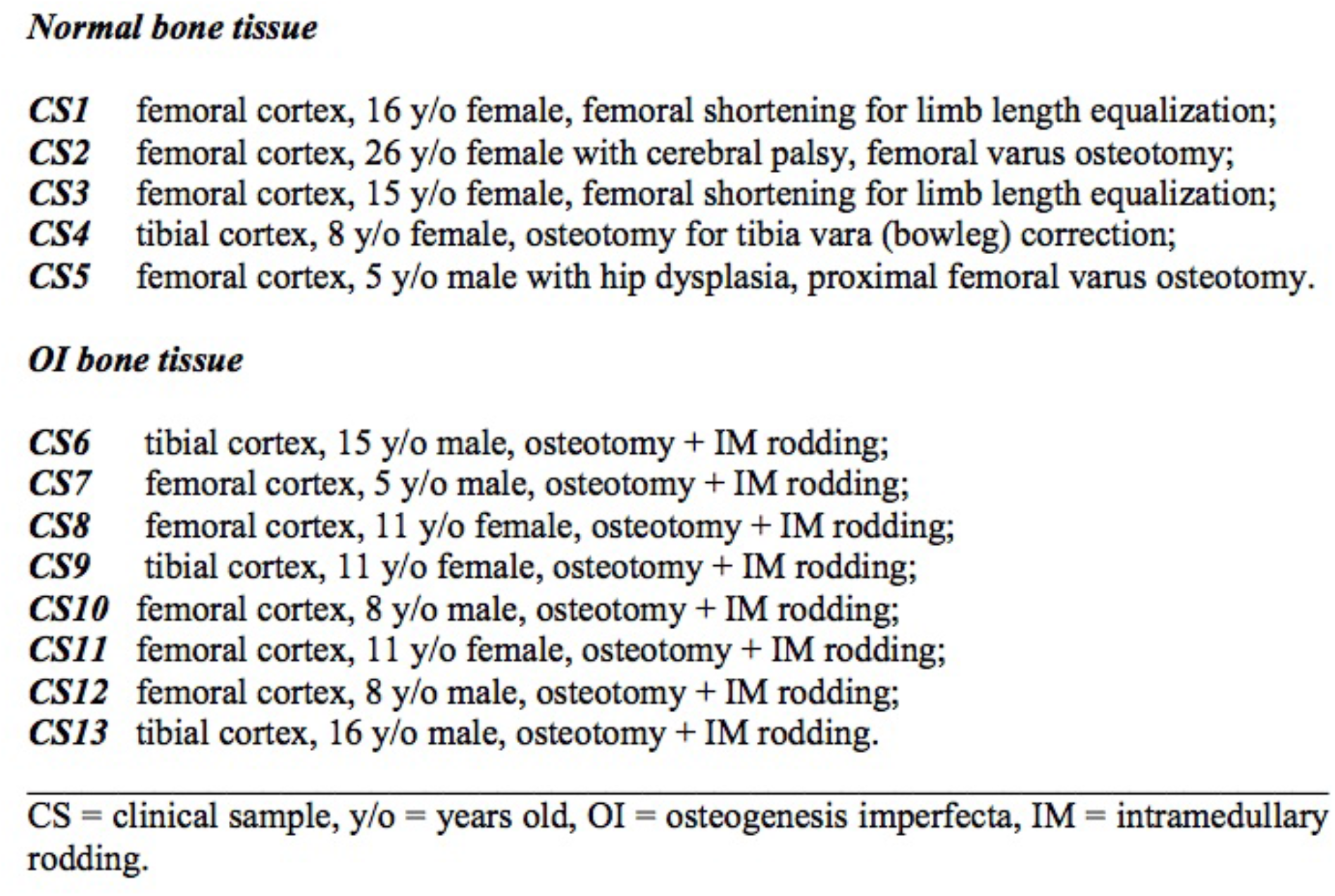
The source of tissue samples obtained for study is listed for each of the normal and OI samples.

### b) Tissue preparation for light and polarization light microscopy

Bone was prepared for microscopy examination using the JB4 plastic technique as follows: bone tissues were fixed in 10% neutral buffered formalin for 3 weeks, decalcified until soft in 25% formic acid, cut into segments approximately 15 mm X 7mm X 7mm or smaller (since larger pieces risk incomplete infiltration by the JB4 medium), infiltrated in JB4 medium (Polysciences, Warrington, PA) for several weeks and subsequently embedded in JB4 plastic. Tissues were then sectioned at 5μm thickness, using a Microm HM 350 rotary microtome (Microm International GmbH, Walldorf, Germany) and stained with 1% toluidine blue.

### c) Tissue examination

*i) Light microscopy (LM)*. LM assessment was performed on an Olympus BX50 photomicroscope equipped for polarization studies, histomorphometric measurements (cellSens imaging software) and digital photography (Olympus CMOS SC 30 digital camera). Assessments included: osteoblast and osteocyte cellularity in relation to matrix structure (woven or lamellar), identification of elongated elliptical “flattened” osteocytes lying along the long axis of the lamellar tissue, orientation of the osteocyte canaliculi, and structure of lamellar tissue deposition. Multiple tissue samples were examined. *ii) Polarizing light microscopy (PLM)*. PLM assessment was then performed with polarization adaptation using a U-AN360P rotatable analyzer and a γ 137 nm U-TP137 polarizing quarter wave mica plate fixed compensator. For demonstration and documentation of the polarization of any specific matrix, a light microscopy section photomicrograph was taken followed immediately by a polarization view of the same section. Figures 1a – 1c illustrate LM and PLM features of normal and OI bone that enable accurate lamellar measurements.

**Figures 1a (i, ii).**
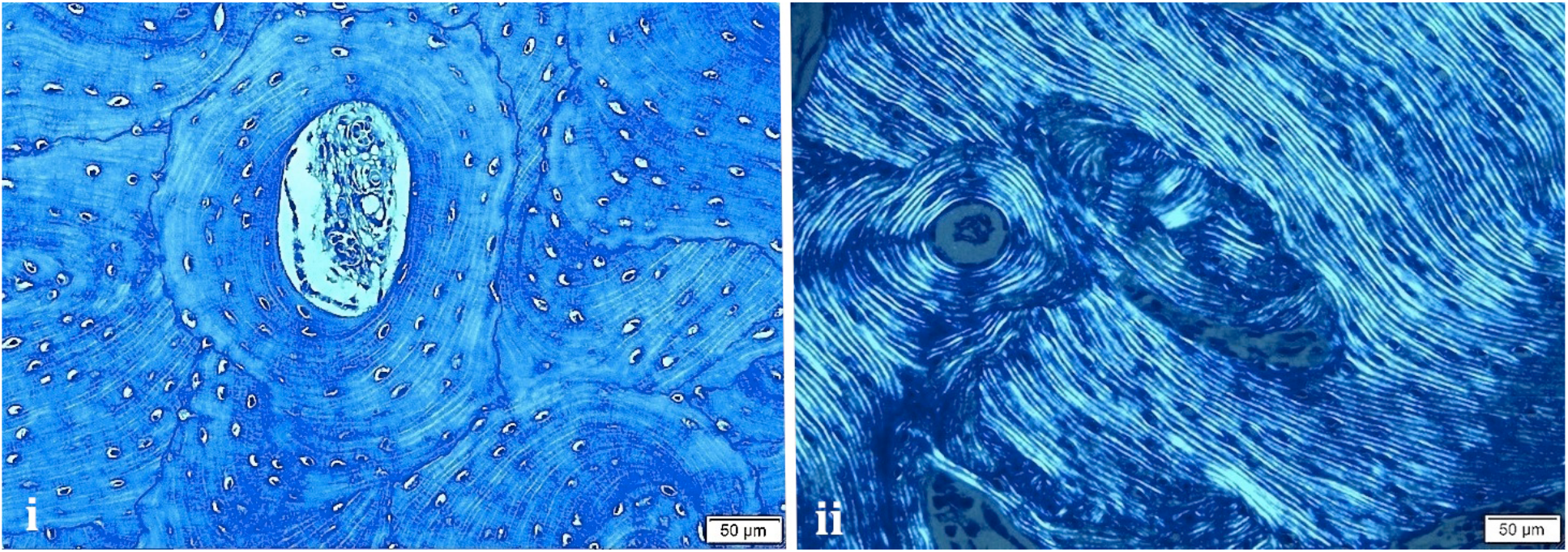
Bone from OI patient showing mosaic lamellar bone tissue with fragments in random positioning. Note central osteon cut in cross-section surrounded by bone oriented in multiple planes but still showing lamellation. i) light microscopic view; ii) polarization microscopy view of adjacent different tissue section demonstrates lamellar structure in all segments more clearly.

**Figures 1b (i – iv).**
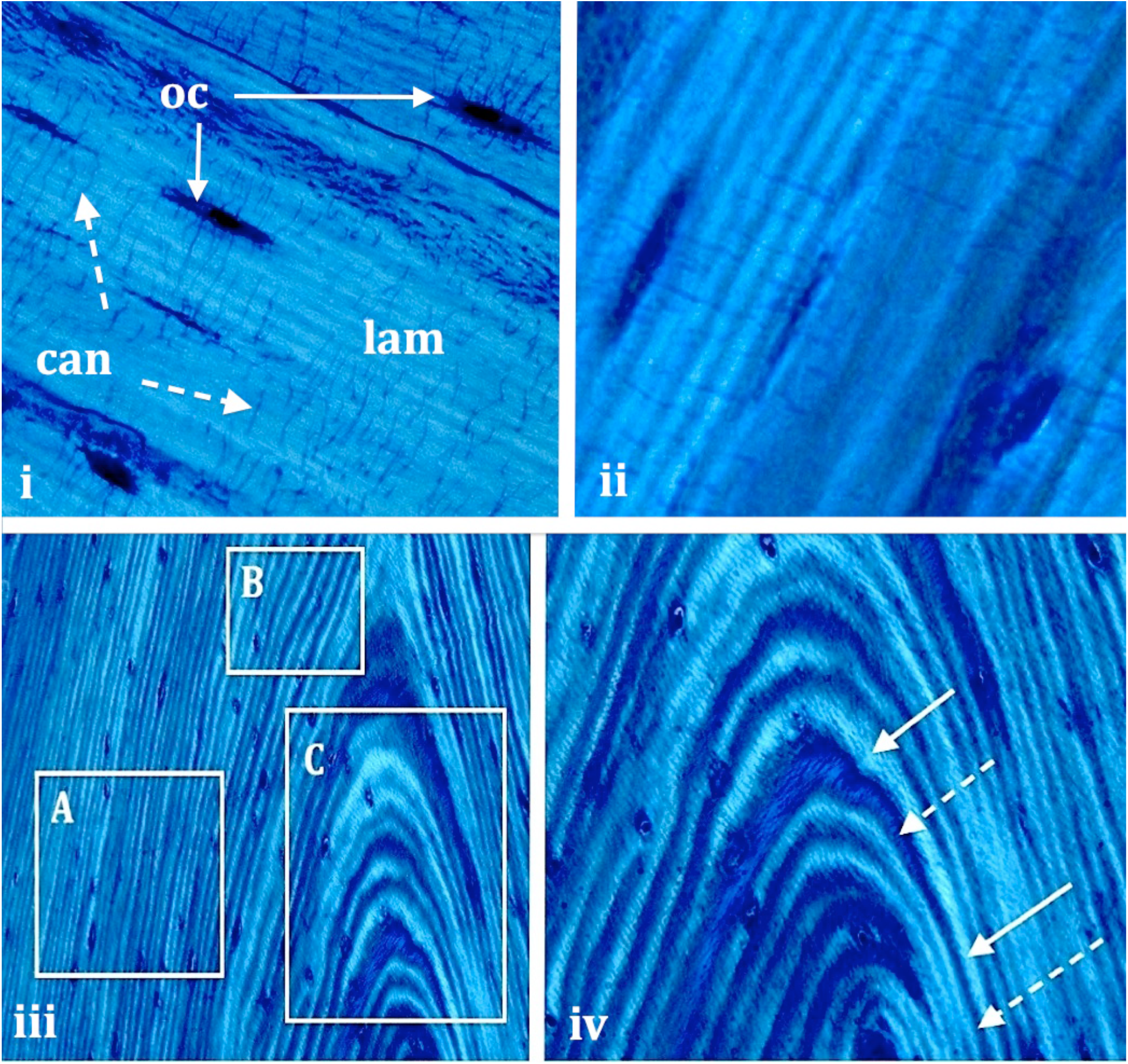
Photomicrographs from cortical bone from normal patient demonstrate the criteria used in choosing regions to measure (or not measure) for lamellar quantitation. Images are shown under “partial” polarization to illustrate both lamellation and cells and canaliculi. i) Lamellae (lam) are visible at center right. Osteocytes (oc) are of flattened elliptical shape and lie along the long axis of the lamellae. Canaliculi (can) are continuous with the oc sidewalls and lie perpendicular to the lamellae, crossing them generally at right angles. Regions such as this are where mature regularly oriented lamellae can be measured. ii) Features seen in i) are shown at higher magnification from different area. iii) Section of normal cortex shows 3 collections of lamellae: box A has uniformly sectioned unidirectional lamellae where measurements can be made; box C has obliquely sectioned lamellae where lamellar thickness varies along the pathway of an individual lamella; areas such as this provide highly inaccurate indications of lamellar thickness and should not be measured; box B shows the beginnings of obliquity; such regions must be carefully assessed and also not measured. iv) Oblique regional view shows thickness changes along individual lamellae, one lamella shown with 2 solid arrows and another with 2 interrupted arrows.

**Figures 1c (i – iii).**
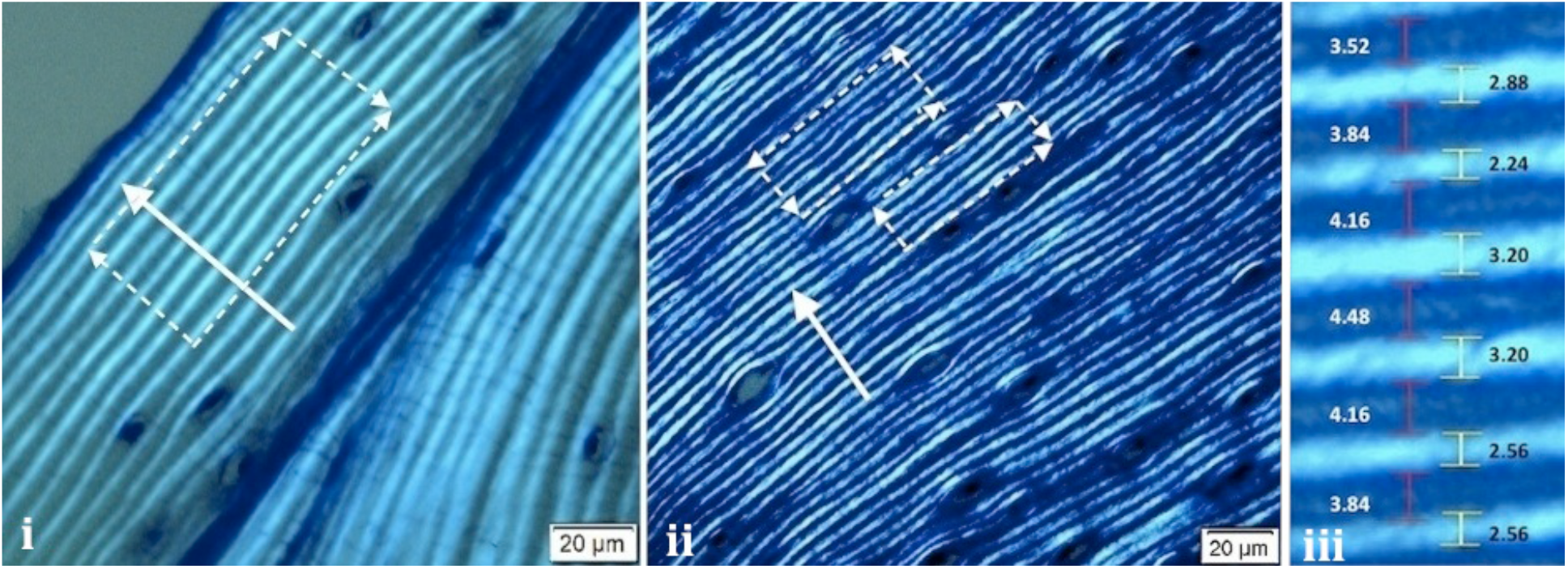
Measurement approaches for normal bone lamellae (i) and OI bone lamellae (ii) under PLM are shown. (i) Normal lamellae show alternating bright and dark bands. Osteocytes are positioned within the wider dark lamellae. The immediately adjacent bright lamella will often be slightly curved and flattened as it ‘curves around’ the osteocyte. For this reason the layers of lamellae should not be measured immediately adjacent to osteocytes. Single arrow delineates an area chosen for the manual measurements. Rectangular area delineates an area chosen for automated thickness averaging. ii) Lamellae in OI bone also show alternating bright and dark bands. The bands are qualitatively noted to be thinner than normal (both images taken at same magnification, 20μm marker). The osteocytes in OI are relatively larger and more oval in shape than those in normal bone. Combined with narrower lamellae, the osteocytes further disrupt the pathway of the lamellae than in normal bone leading to a more ‘wavy’ appearance of the lamellae and often to discontinuous lamellae. iii) High power photomicrograph of a region under PLM that was measured manually is shown from a screenshot of an actual measurement. The thickness of individual bright and dark lamellae as measured by histomorphometry is shown. Lamellar thickness of dark lamellae in micrometers is shown at left (white numbers, pale red marker) and the bright lamellae at right (yellow marker, black numbering).

### d) Measurement of lamellar thicknesses

#### i) Choosing which lamellar group concentrations to measure

Bone sections were examined using LM and PLM, at lower power 4X and 10X magnifications, to choose 3 to 5 areas per section for measurement. Criteria for measurement included: 1) relatively large accumulations of 16-20 bright and dark lamellae with 2) qualitatively regular thicknesses throughout, 3) flattened elliptical mature osteocytes along the longitudinal axis of the lamellae, 4) canaliculi passing from the walls of the osteocyte lacunae at right angles to the lamellae, and 5) (in cross-sections of bone) circular osteons cut without obliquity as indicated by a uniform number of lamellae circumferentially arrayed around the central circular vascular space. These criteria were designed to eliminate oblique lamellae from quantitation, since any obliquity would lengthen the thicknesses measured (Figure 1 a - c). Once regions for quantitation were chosen examination with PLM was done at 20X magnification. The microscope stage was rotated with both polarizers positioned to obtain the sharpest delineation of dark and light lamellae. Care was taken to focus on the bright lamellae so that their edges were as sharp as possible since blurred edges falsely widened the thicknesses. Lamellae sectioned along the longitudinal axis were more commonly measured but some lamellae in cross-section (in classically recognized osteons) were also quantitated. The regions chosen for histomorphometric quantitation were photographed and labeled allowing for measurements to be done from the captured images.

#### ii) Histomorphometry of lamellae

Bright and dark lamellar thicknesses were measured in regions chosen for quantitation using the Olympus microscope cellSens image analysis histomorphometry software system. We refer to this approach as the manual method since, using the cellSens imaging software, a cursor marking is manually positioned to outline the extent of each lamella and determine its thickness in micrometers (Figure 1c). Accuracy and ease of measurement were improved at the 20X magnification by increasing the localized magnification to 150% - 200% that still maintained sharp differentiation between the extents of bright and dark lamellae. Magnification beyond 200% tended to blur the borders between the layers.

#### iii) Development of automated thickness averaging method to ease quantitation

A script for automated measurement of average lamellar thicknesses from microscopic images was developed in MATLAB (Mathworks, Natick, MA). The script measured the thickness of each lamella contained within a rectangular measurement area by averaging multiple measurements across the length of the area (compared with a single measurement for each lamella using the manual method). The steps involved in the automated thickness averaging method are shown in figures 2 a – d. After the measurement region was manually selected and marked (Figure 2a), the region was automatically rotated so that the lamellae were oriented horizontally. Contrast was enhanced by setting the intensity of the darkest and brightest 1% of pixels to the minimum and maximum intensity values, respectively. Remaining pixels were assigned new intensities based on their initial intensities relative to the minimum and maximum (Figure 2b). The image was binarized using a local thresholding (average of 20 surrounding pixels in column). Pixels with intensity higher than the local average were marked as belonging to bright lamellae, and pixels with lower intensity were marked as dark lamellae (Figure 2c). ‘Floating’ pixels were recolored to fit the lamella they are found in. Columns containing edge artefacts or the wrong number of lamellae were removed from measurement (Figure 2d). Lamellar thickness was measured across the length of the entire measurement window, with a measurement of each lamella being taken in each column of pixels. Pixels were square with side length 0.3185 μm. The average of all measurements for a single lamella was taken to produce the final value for its thickness. All measurement areas contained a consistent number of lamellae across the length of the image. The mean number of measurements for each lamella was 86 for normal bone and 72 for OI bone. The automated thickness averaging method was used to measure lamellae in bone along both the transverse and longitudinal axes.

**Figures 2a-2d.**
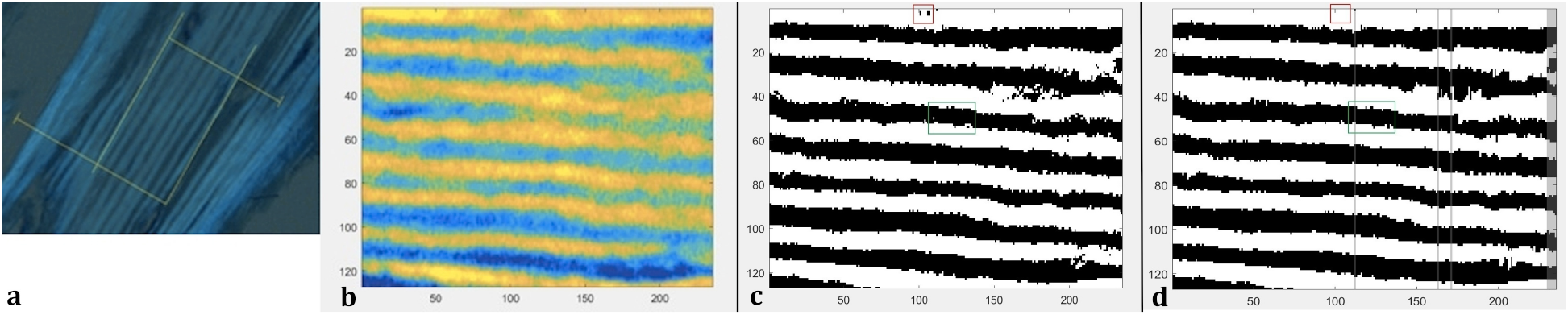
Images of the measurement of a normal bone sample (CS4) throughout several steps of image processing are shown. a) An image of the sample as viewed with polarized light at 20X magnification is shown. The rectangle marking the measurement area has been created using two sets of perpendicular lines. b) The measurement area is seen after it has been cropped and reoriented so that the lamellae run horizontally. Contrast has been boosted to differentiate between bright and dark more clearly. c) The measurement area after each pixel has been marked either bright or dark relative to the local average intensity is shown. Boxed regions contain floating pixels that will be altered to match their surroundings. d) The measurement area is seen after floating pixels have been corrected. Columns excluded from measurement have been grayed out.

#### iv) Validation of automated thickness averaging method

To confirm consistency between manual and automatic measurements, a validation dataset was created. The same samples of normal and OI bone measured by the automated averaging script were subsequently measured by the manual method and these measurements were compared against one another. This dataset includes measurements of 423 lamellae, 211 lamellae from three normal patients (without any molecular abnormalities) and 212 lamellae from five patients with OI.

### e) Measurements made in study

#### i) Manual measurements alone

Manual measurements were made in patients listed in Table 1 involving all 5 of the normal tissue samples (C1 – C5) and 7 of the OI tissue samples (C6 – C12). This group involved 1811 lamellae, 707 lamellae from the 5 normal patients and 1104 lamellae from the 7 OI patients. *ii) Automated thickness averaging method and its validation comparing consistency between automated and manual measurements*. The tissue samples assessed for the automated thickness averaging method were chosen from selected patients listed in table 1 but different histologic slides were used such that different regions of lamellae were measured from those measured solely by the manual method described in i) above. This group involved 423 lamellae, 211 lamellae from 3 of the normal patients (CS1, CS2 and CS4) and 212 lamellae from 5 of the OI patients (CS6, CS7, CS8, CS11 and CS13).

### f. Statistical analyses used to assess measurements made

Statistical analyses were performed separately for the lamellae assessed by manual measurements alone (section e (i) above, 1811 lamellae) and for the different set of lamellae assessed by the automated thickness averaging method and then validated for consistency by subsequent measurements (on the same lamellae) using the manual method (section e (ii) above, 423 lamellae).

#### i) Statistical analyses used to assess manual measurements alone (1811 lamellae)

A two-sided ANOVA followed by a multiple comparison test with Tukey-HSD post hoc was applied to determine statistical significance in lamellar thickness for patient group (normal or OI) and for lamellar type (bright or dark) (α = 0.05).

#### ii) Statistical analyses used to assess consistency between automated and manual measurements (423 lamellae)

Paired statistical tests were used to determine whether the measurement made by the automated averaging method differed significantly from the manual measurements on the same lamellae. A Lilliefors test determined that the differences between the automated and manual measurements of the same lamellae were not normally distributed (α = 0.05). For this reason, a non-parametric test was used. The equivalence of automated and manual measurements of the same lamellae was evaluated using a two-sided sign test with the null hypothesis that the median difference between the measurements was 0 (α = 0.05).

## RESULTS

### a) Qualitative description of appearances of normal and osteogenesis imperfecta lamellae

Normal bone is characterized by the presence of uniform, well-defined lamellar bands (Figure 1c (i)). Lamellae in OI bone tend to be slightly less well defined and more irregular than their counterparts in normal bone. The lamellar bone in OI is more cellular and the osteocytes are more oval than cells in normal bone as seen in Figures 1a (ii) and 1c (ii). In OI, the combination of more cells, relatively larger cells and oval shaped cells (in comparison with flattened elliptical osteocytes in normal lamellar bone) renders associated lamellae ‘wavy’ in conformation. This occurs as lamellae ‘curve around’ adjacent osteocytes. Also, the relatively large and oval osteocytes in OI bone can simply interrupt lamellae that are normally continuous around the cells. These characteristics often limited the size of potential measurement areas in slides from OI bone. Areas where two bright or dark lamellae blurred together were more common in OI bone and these had to be excluded because the script could not identify the boundary between them. Areas containing large, rounded cells were also excluded from measurement because they appeared to the counting algorithm as large sections of dark lamellae. OI lamellar bands tended to organize less uniformly, often merging and diverging with adjacent lamellae and creating spindle-shaped layers that make OI lamellae less sharply defined. In comparison to normal bone lamellae, instances of measurable OI lamellae were less frequent. The decreased number of clearly defined areas due to non-uniformity, immaturity, and increased cellularity presented a challenge to lamellae quantitation. For these reasons, measurement areas in OI bone samples were often slightly smaller than in normal bone samples.

### b) Quantitative histomorphometric measurements of normal and osteogenesis imperfecta bone bright and dark lamellar thicknesses

The values measured for the lamellar thicknesses (group of 1812 lamellae) are shown in figures 3 a – c and table 2 below. The distributions of thicknesses from bright and dark lamellae in normal and OI bone are shown in Figures 4 a and b.

**Figure 3a.**
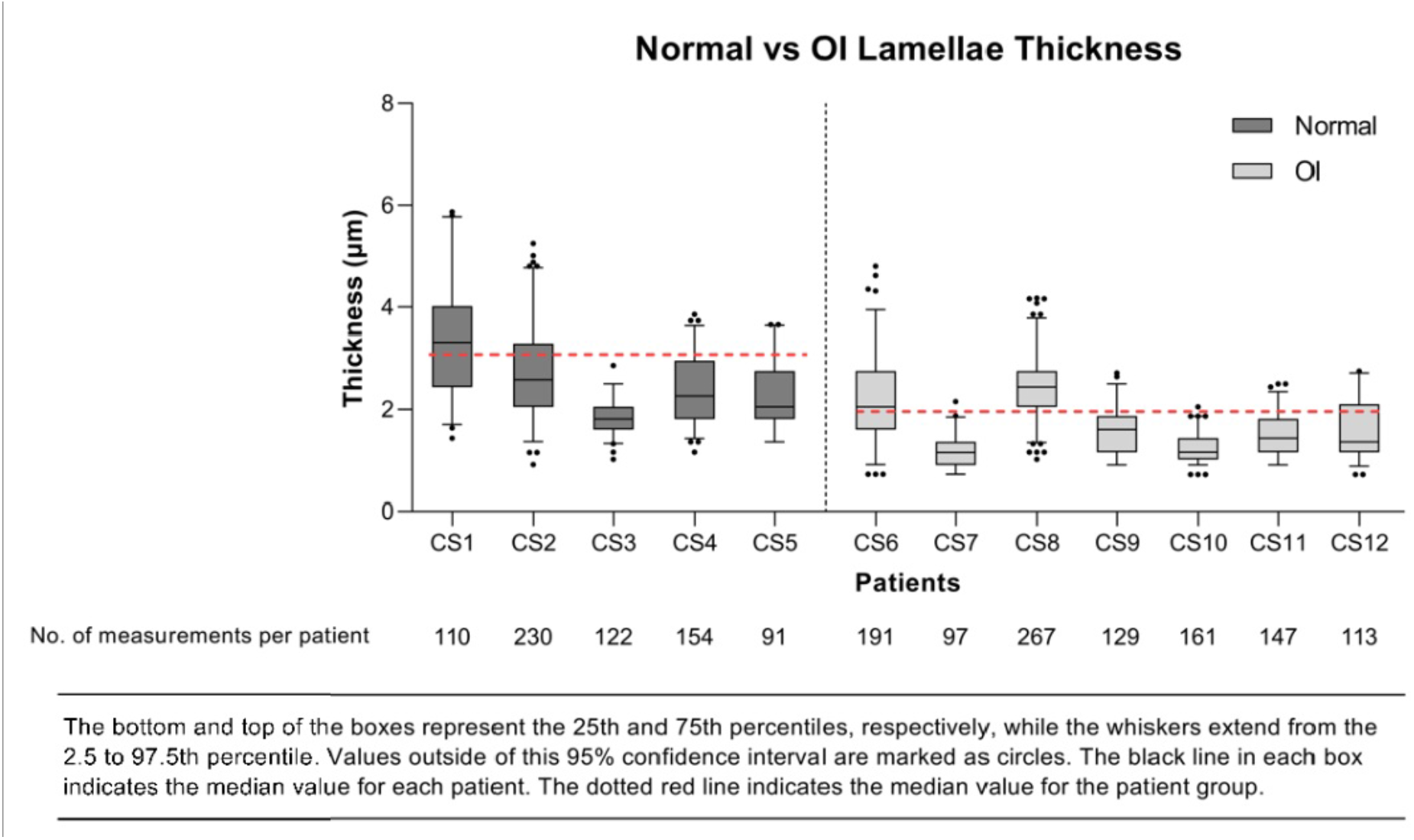
The normal vs OI lamellar thicknesses are shown. This measurement is based on the median value for bright and dark lamellar thicknesses combined for each patient.

**Figure 3b.**
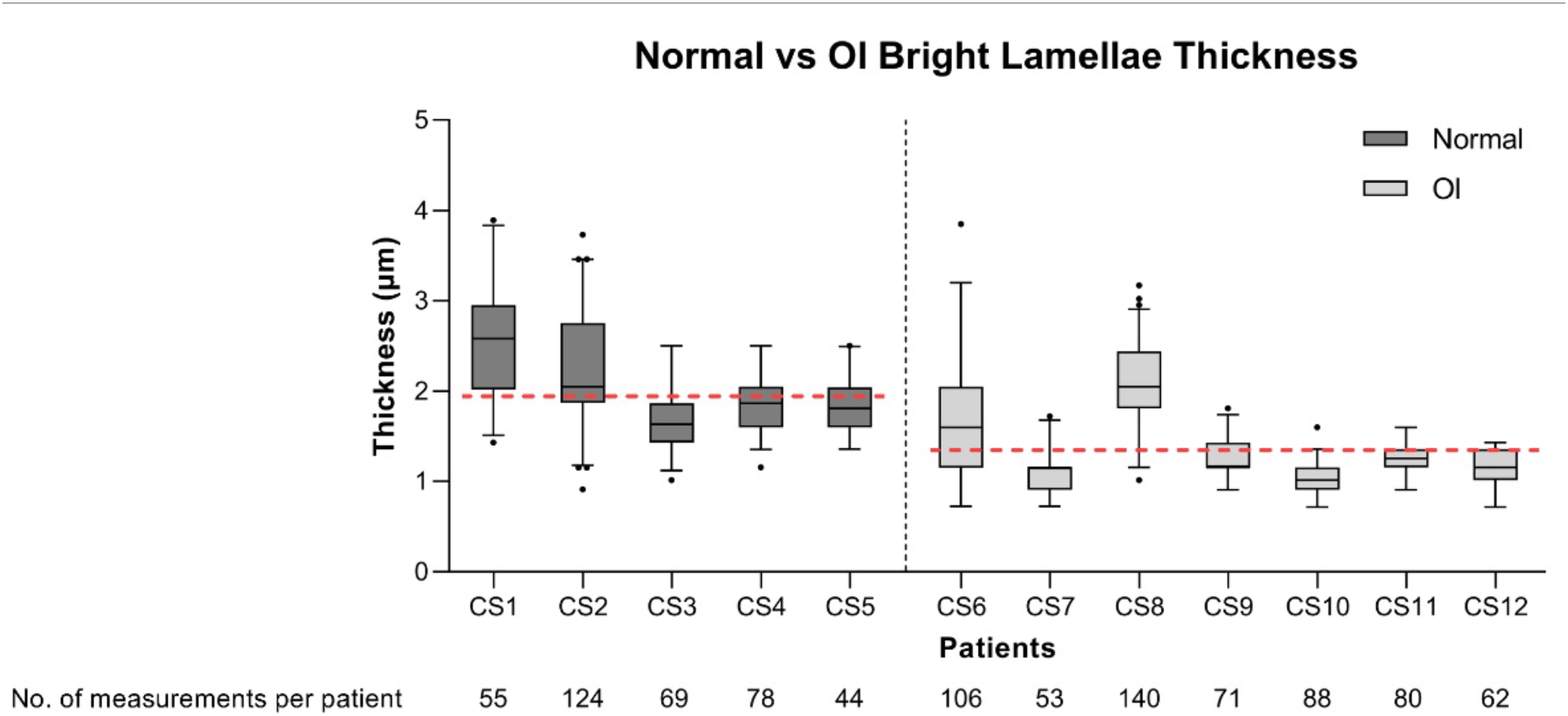
The thicknesses for the bright lamellae are listed for normal and OI patients.

**Figure 3c.**
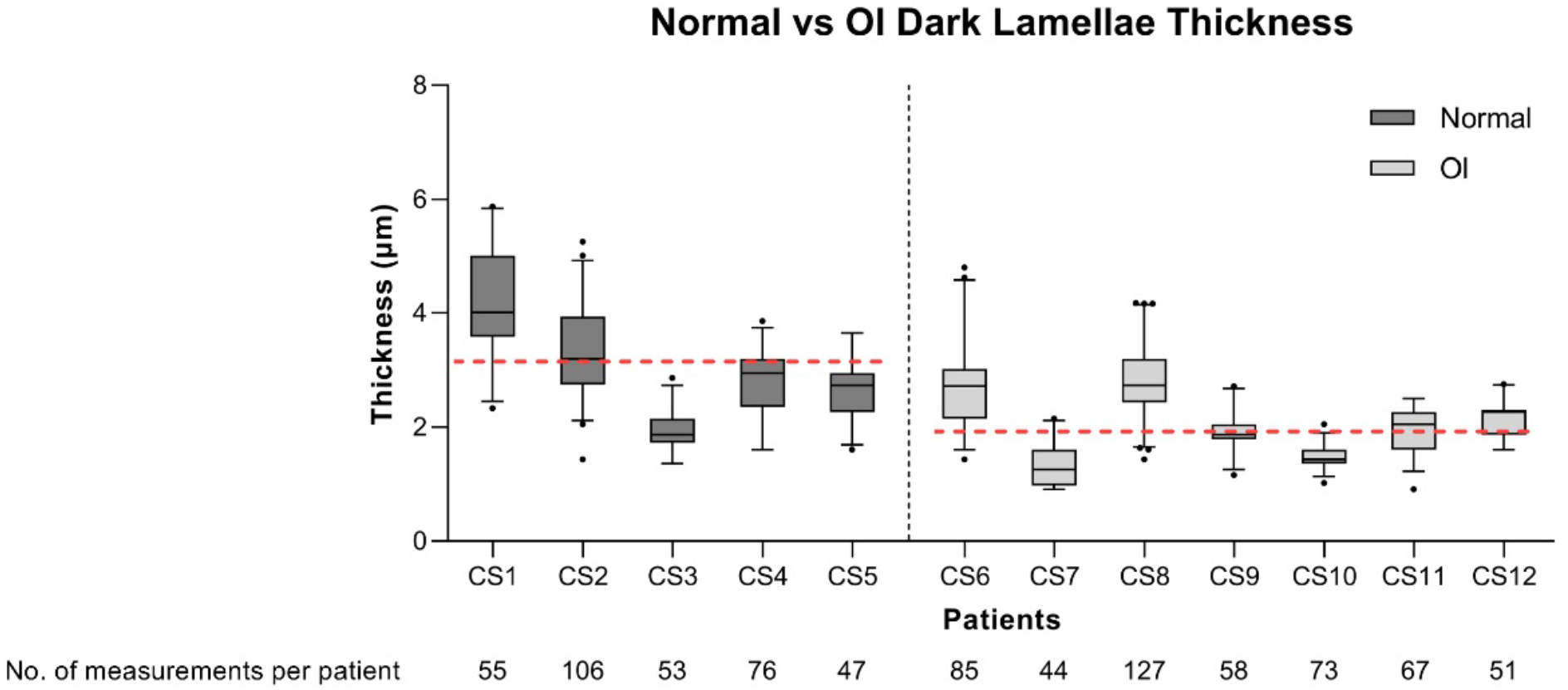
The thicknesses for the dark lamellae are listed for normal and OI patients.

**Table 2.**
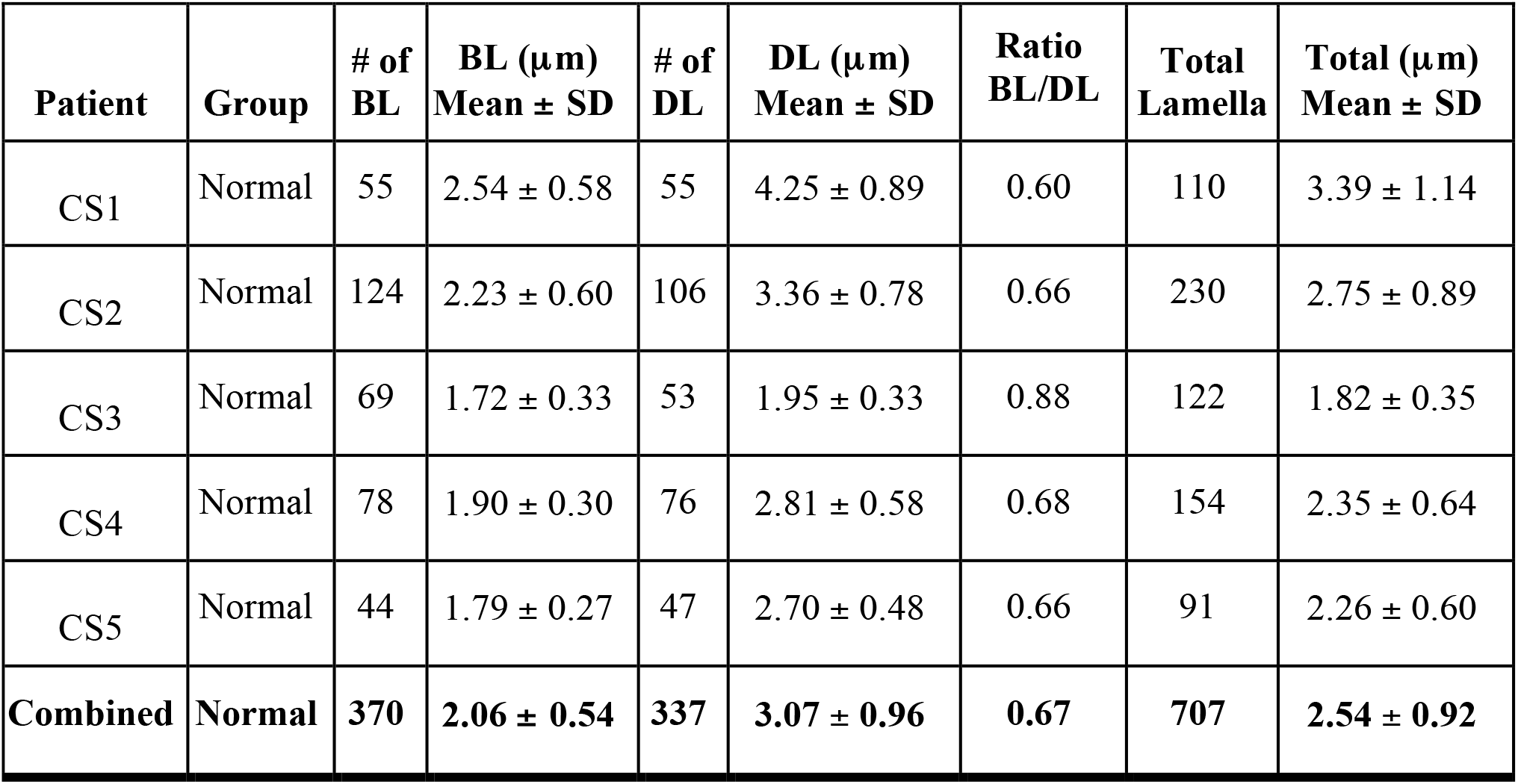
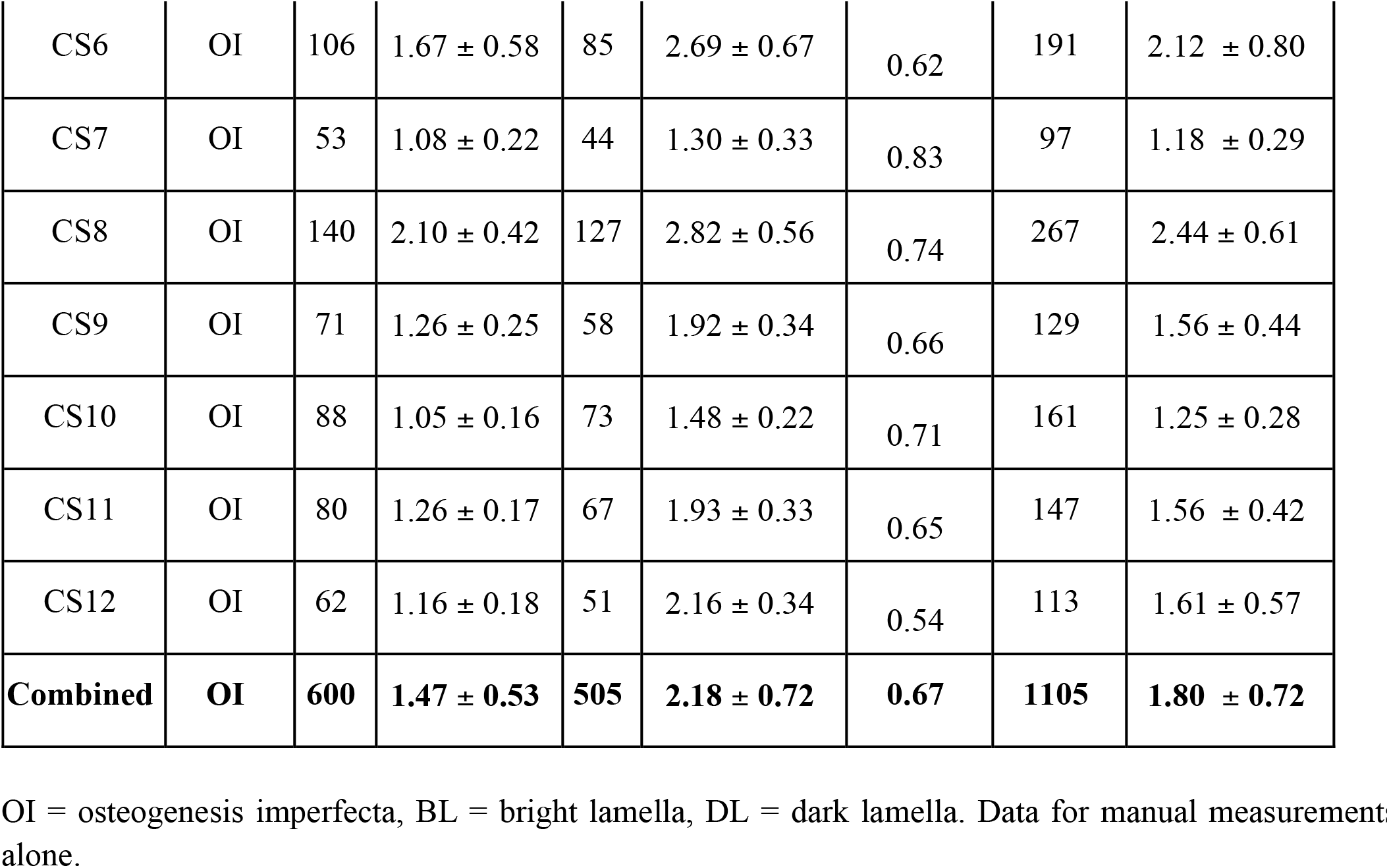
Bright lamellar and dark lamellar thicknesses and ratios in normal and osteogenesis imperfecta bone.

**Figures 4 (left) and 4b (right).**
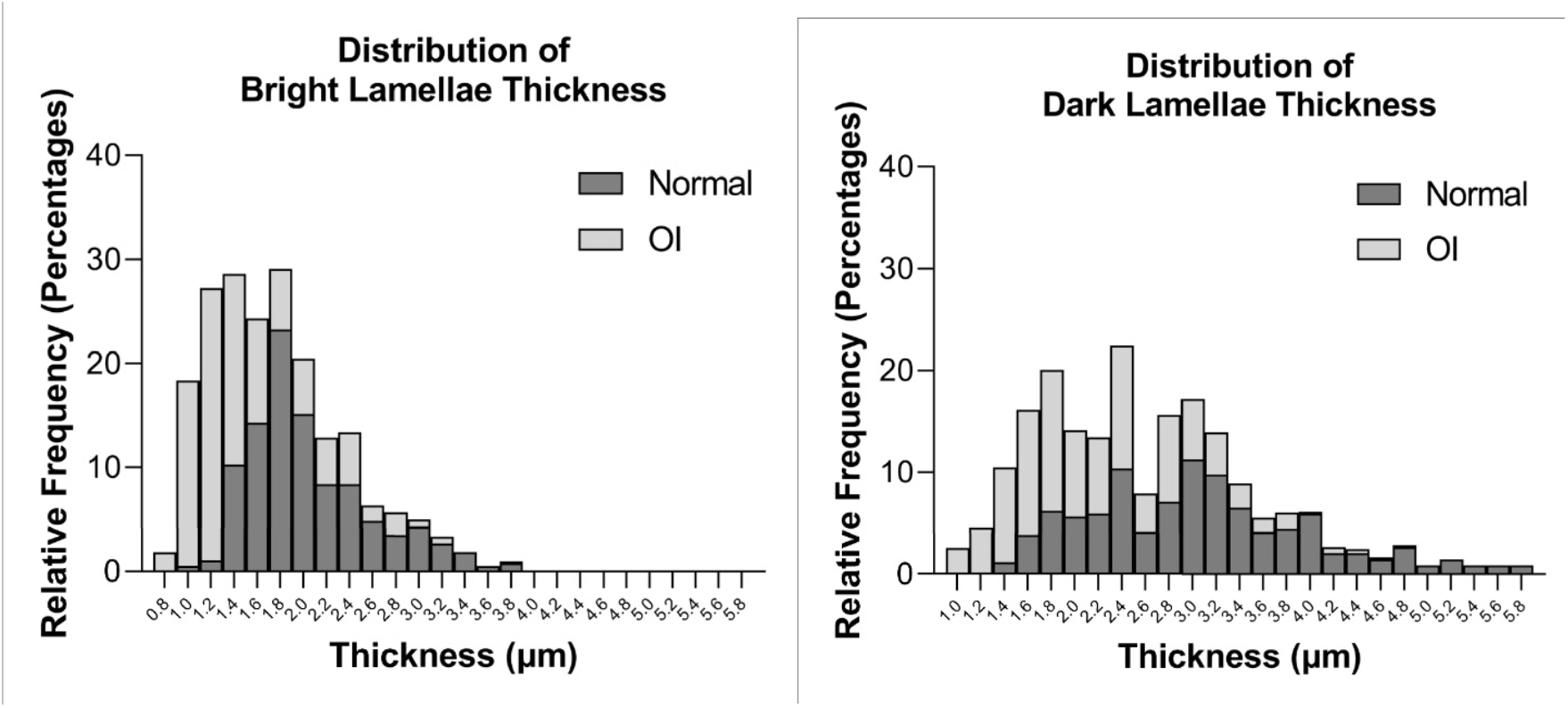
The thickness distributions for bright lamellae in normal and OI bone are illustrated at left (4a) and for dark lamellae in normal and OI bone at right (4b).

#### Statistical comparison of groups

A two-way ANOVA on the manual measurements alone group defined a statistically significant difference for patient group (f(1) = 496.06, p<0.001) and for lamellar type (f(1) = 675.54, p<0.001), with a statistically significant interaction between these factors (f(1) = 19.85, p<0.001).

A multiple comparison test with post hoc analysis using a Tukey-HSD test determined statistically significant differences in the average lamellar thickness between normal bright lamellae and OI bright lamellae (p<0.001), normal dark lamellae and OI dark lamellae (p<0.001), normal bright lamellae and normal dark lamellae (p<0.001), and OI bright lamellae and OI dark lamellae (p<0.001). In comparisons between patient groups, OI bright and dark lamellae were found to be thinner than normal bright and dark lamellae respectively. In comparisons between lamellar type within a patient group, bright lamellae were found to be thinner than dark lamellae.

### c) Validation dataset to compare the percent difference between the manual and automated measurements

For each lamella in the validation dataset, the percent difference between automated and manual measurements was calculated. This comparative validation involves automated thickness measurements and manual measurements made on the same lamellae. The data for this validation assessment that included 423 lamellae are shown in Table 3 and Figures 5a and b. The mean of the absolute values of these percent differences was 18.9%.

**Table 3.**
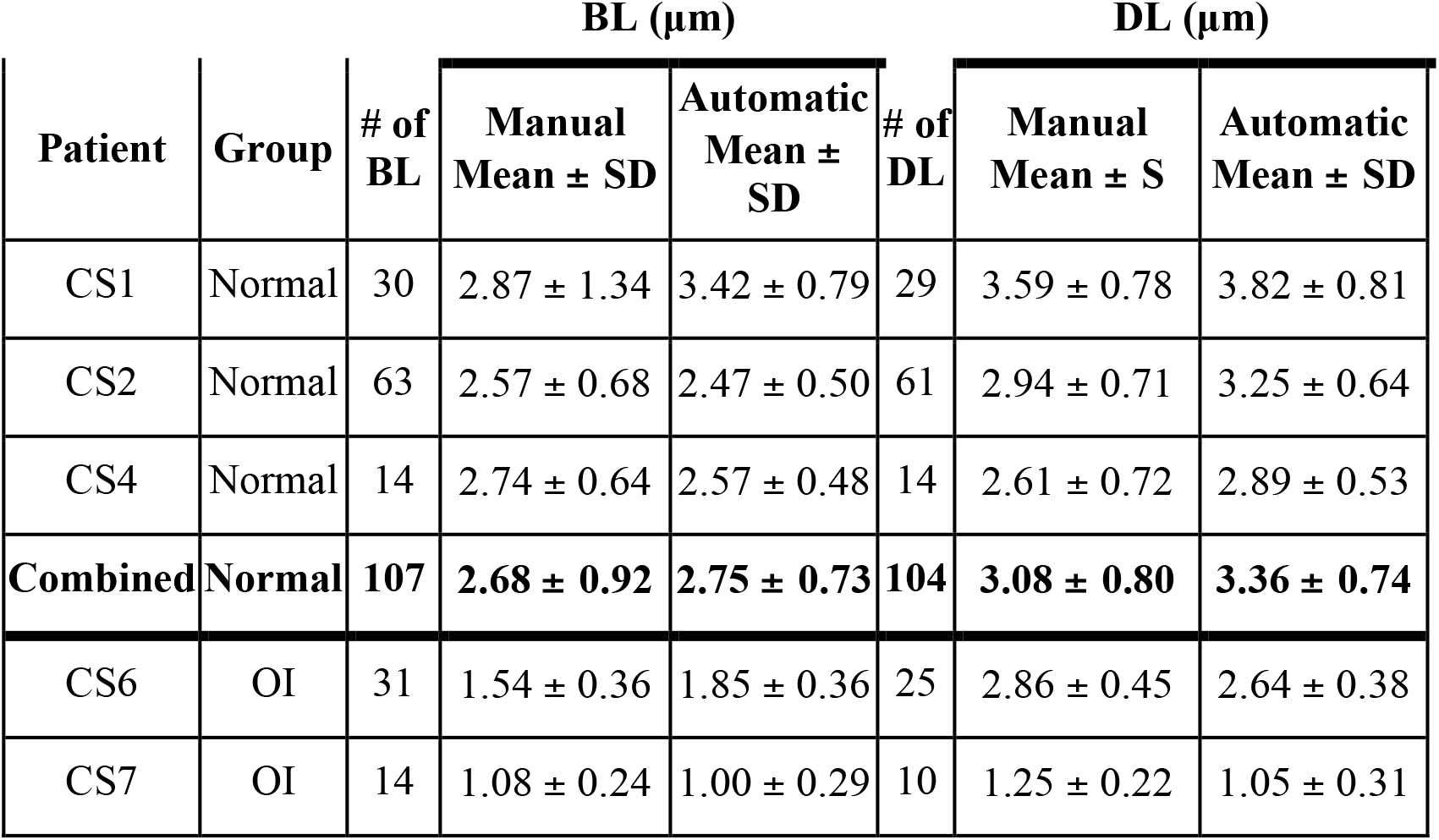
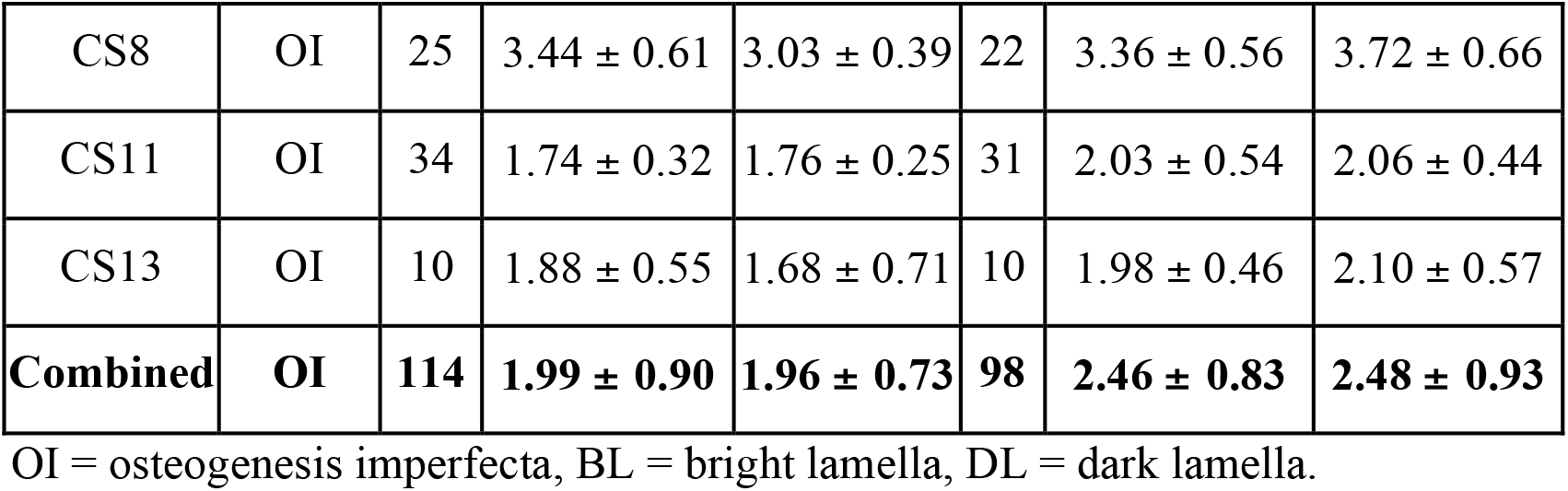
Data indicate comparison of automatic and manual methods for comparing thicknesses of bright and dark lamellae.

| Patient | Group | # of BL | BL ($\mu\text{m}$) | | # of DL | DL ($\mu\text{m}$) | |
| --- | --- | --- | --- | --- | --- | --- | --- |
| | | | Manual Mean ± SD | Automatic Mean ± SD | | Manual Mean ± S | Automatic Mean ± SD |
| CS1 | Normal | 30 | $2.87 \pm 1.34$ | $3.42 \pm 0.79$ | 29 | $3.59 \pm 0.78$ | $3.82 \pm 0.81$ |
| CS2 | Normal | 63 | $2.57 \pm 0.68$ | $2.47 \pm 0.50$ | 61 | $2.94 \pm 0.71$ | $3.25 \pm 0.64$ |
| CS4 | Normal | 14 | $2.74 \pm 0.64$ | $2.57 \pm 0.48$ | 14 | $2.61 \pm 0.72$ | $2.89 \pm 0.53$ |
| **Combined** | **Normal** | **107** | **$2.68 \pm 0.92$** | **$2.75 \pm 0.73$** | **104** | **$3.08 \pm 0.80$** | **$3.36 \pm 0.74$** |
| CS6 | OI | 31 | $1.54 \pm 0.36$ | $1.85 \pm 0.36$ | 25 | $2.86 \pm 0.45$ | $2.64 \pm 0.38$ |
| CS7 | OI | 14 | $1.08 \pm 0.24$ | $1.00 \pm 0.29$ | 10 | $1.25 \pm 0.22$ | $1.05 \pm 0.31$ |
| CS8 | OI | 25 | $3.44 \pm 0.61$ | $3.03 \pm 0.39$ | 22 | $3.36 \pm 0.56$ | $3.72 \pm 0.66$ |
| CS11 | OI | 34 | $1.74 \pm 0.32$ | $1.76 \pm 0.25$ | 31 | $2.03 \pm 0.54$ | $2.06 \pm 0.44$ |
| CS13 | OI | 10 | $1.88 \pm 0.55$ | $1.68 \pm 0.71$ | 10 | $1.98 \pm 0.46$ | $2.10 \pm 0.57$ |
| **Combined** | **OI** | **114** | **$1.99 \pm 0.90$** | **$1.96 \pm 0.73$** | **98** | **$2.46 \pm 0.83$** | **$2.48 \pm 0.93$** |

OI = osteogenesis imperfecta, BL = bright lamella, DL = dark lamella.

**Figure 5a.**
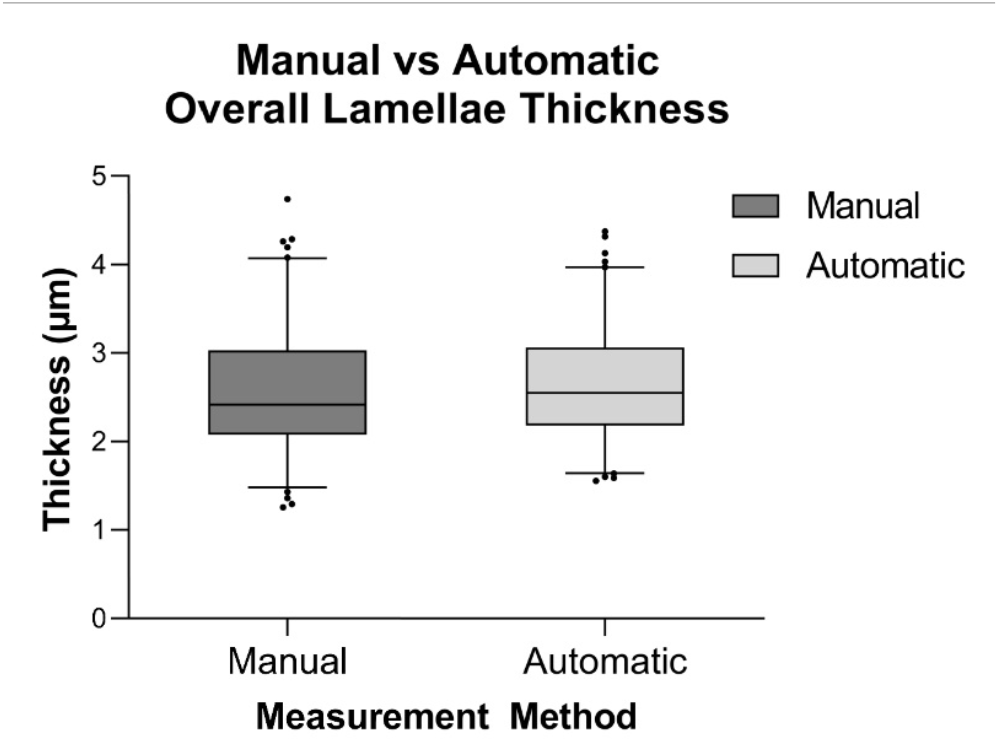
The overall lamellar thickness obtained using manual measurements and automated thickness averaging method measurements are shown.

**Figure 5b.**
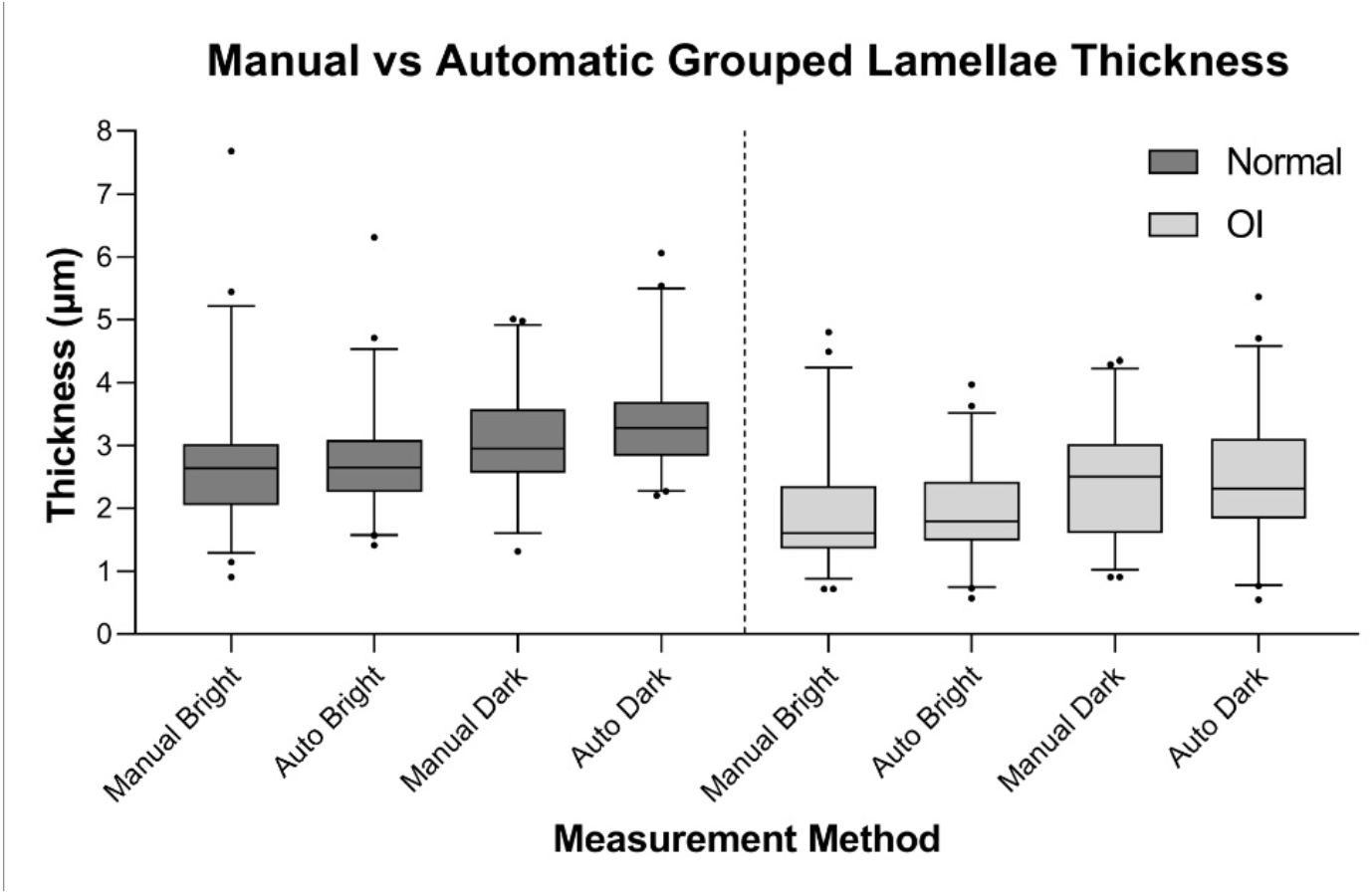
The manual compared with automatic method measurements for each specific group are shown.

There were several lamellae for which the automatically measured thickness was much greater than the manually measured thickness; in rare cases, the automated measurement was more than twice as thick. Closer examination of these measurements showed that this occurred when the manual measurement of a lamella was taken at its thinnest point (Figure 6).

**Figure 6.**
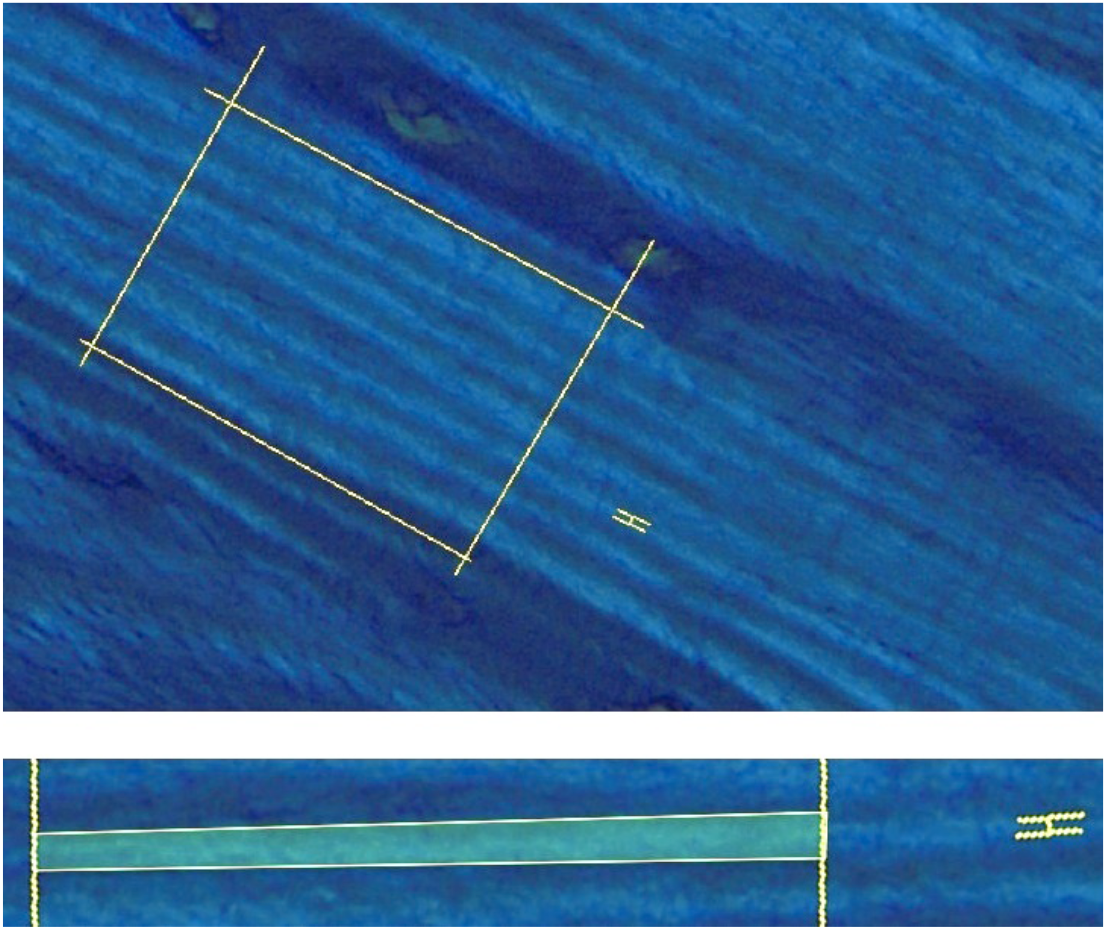
An examination of the lamella with the largest discrepancy between automated and manual measurements is shown. The top image shows the positions of the rectangular automated measurement area and the manual measurement of the lamella in question. The yellow marker at right shows how the lamella under consideration is measured manually at one position. The bottom image, rotated and zoomed in, shows that the lamella is thinner at the location of manual measurement and thicker inside the entire automated measurement area. This lamella had a manual measurement of 1.43μm and an automated measurement of 3.09μm, a difference of 116.1%.

A Lilliefors test determined that the differences between the automated and manual measurements of the same lamellae were not normally distributed (p<0.005). A two-sided paired sign test on the median of the differences did not show that the automated and manual measurements were significantly different (p = 0.0518).

## DISCUSSION

### Two components of this study outlining advances in the field of bone lamellar interpretation

#### 1. Measurements comparing lamellar thicknesses in OI and normal bone

The data comparing measurements of lamellar thicknesses in OI and normal bone allow for three specific observations: i) The OI bone mean lamellar thickness values (bright and dark lamellae combined) are always less than those in normal bone (OI 1.80 ± 0.72 μm and normal 2.54 ± 0.92 μm). ii) The mean value for the bright lamellae is less than that for the dark lamellae in both normal and OI bone (OI: bright lamellae 1.47 ± 0.53 μm and dark lamellae 2.18 ± 0.72 μm and normal bone: bright lamellae 2.06 ± 0.54 μm and dark lamellae 3.07 ± 0.96 μm). iii) The ratio of mean values for bright/dark lamellar thicknesses is the same in OI and normal bone: OI bone ratio 0.67 (range: 0.54 – 0.83) and normal bone ratio 0.67 (range: 0.60 – 0.88). The histograms of lamellar thicknesses clearly illustrate the difference between OI and normal lamellae. Differences in the thickness of OI bone lamellae compared to normal bone have not previously been recognized.

#### 2. Development of a method for automated thickness averaging measurement of lamellae thicknesses

Since measurement of the lamellar thicknesses in sufficient numbers to provide statistical validity is time consuming, we have developed a method for automated thickness averaging in an effort to measure lamellar thickness more quickly. This will allow for lamellar measurements to become more widely used in research and as a clinical laboratory tool enhancing histomorphometric assessments of normal and pathologic bone. Automated and manual measurements of the same lamellae in the validation dataset were shown not to be significantly different. Automated lamellar thickness measurements provide a more robust assessment of lamellar thickness, as averaging thickness over a wide area accounts for varying thickness of a single lamella better than a single-point measurement. Additionally, this automated method removes some of the subjectivity of manual measurements.

### Previous assessments of lamellar thicknesses in bone

#### 1. Lamellar thicknesses in human bone

Lamellae in cortical bone have been clearly defined since the mid 1800s and polarization microscopy was invented almost 200 years ago. Although lamellae are integral to bone structure, lamellar thicknesses in normal human bone have been reported infrequently, with a relatively wide range of values, and have not been used in clinical studies while those in OI have not been documented at all. Statements made in general reviews (as distinct from specific research studies) have tended to place the mean lamellar thicknesses in a 3-7μm range (Rho et al, 1998; Jee, 2001). Reports where the lamellar thicknesses have been specifically assessed show the following findings. Frost (1962) reported relatively wide values in his study of human bone lamellae from 41 patients. The study was done using undecalcified fuchsin-stained 50 μm thick sections using polarized light with measurements by an ocular micrometer. A mean thickness was documented of 7.27 μm in outer circumferential lamellae and 7.54 μm in Haversian lamellae. Assessments done in multiple bones (rib, femur and clavicle most commonly) with ages ranging from fetal to 79 years showed extremely small age variation; Haversian lamellae in normal bone from those under 20 years of age averaged 7.25 μm thickness and in those over 20 years 7.78 μm. The method of measurement was not clearly illustrated; however, the most detailed descriptive phrase indicated that it comprised “the distance between the middle of one bright band to its immediate neighboring bright band” which implies that a lamella was considered to be composed of one dark and one bright component. The mean for a single lamella would thus be one half of the value reported, a number more consistent with measurements subsequently reported. Kragstrup et al (1983) used undecalcified 7μm thick sections stained with Masson trichrome examined by polarizing light microscopy to assess lamellar thicknesses in 64 normal human iliac crest biopsies equally divided between males and females. They assessed trabecular bone, also measured a “double lamella” composed of bright and dark components and calculated a mean thickness of 6.44 ± 0.35 μm (equivalent to an average 3.22 μm thickness per single lamella). Frost (1962) and Kragstrup et al (1983) considered the lamellar thin layer adjacent to a thick layer combined as a “basic repeating unit” of lamellar bone that explains measuring them together although the layers appear to represent part of a continuous pattern rather than an isolatable structural entity. Another study in humans, primarily using SEM, assessed normal femoral osteonal bone from 7 patients 2 months - 43 years of age and showed mean values for lamellar thicknesses of 2.94 ± 0.62 μm (Reid, 1986). Marotti (1993), assessing normal human tibial cortical bone from 4 males 9 – 70 years of age (using both PLM and SEM) showed alternating collagen-rich dense layers and collagen-poor loose layers with the dense layers always thinner and bright (anisotropic) and the loose layers thicker and dark (isotropic). The measured differences were highly statistically significant (p<0.001) with the dense (bright) layers 2.0 μm thick and the loose (dark) layers 3.8 μm thick. Another study from the adult tibia in 3 males and 3 females (21 – 72 years old) assessed cross-sections of undecalcified bone 30 – 40 μm thick using SEM; differences in the 2 types of lamellae were documented with the mean thicknesses of dense lamellae (bright by our study) 1.5 μm and mean thicknesses of loose (dark) lamellae ranging from 3 to 5.5 μm (mean ∼4.5 μm) depending on position in the osteon (Ardizzoni, 2001). Pazzaglia et al (2012) studied bone lamellae from 4 male tibias from patients 25 – 52 years old assessing the same regions by SEM and PLM (40 μm thick sections) but their assessments were directed primarily towards determining the structural components of the layers of transversely sectioned osteons. As part of the study a mean thickness determination of 9.0 ± 2.3 μm was reached. The values ranged from 1.9 μm to 66.3 μm indicating their choice of lamellae differed markedly from what we have measured in this study.

#### 2. Absence of measurement of lamellar thicknesses in OI bone

There have been virtually no studies of OI bone lamellae, however, aside from mention of isolated cases. Frost (1962) reported one OI case in an 11-year old boy from his study; the lamellar thickness reported, 5.3 μm, was the lowest value amongst his 41 patients where the average value was 7.27 μm, a finding consistent with a smaller lamellar thickness in OI bone. Reid and Boyde (1985) reported in an abstract on 2 female OI patients, 7 and 8.5 years old, where OI bone also showed thinner lamellae compared to normal bone although measurements were not provided. They indicated: “polarized light microscopy…showed that the interlamellar distance in both OI cases was half the normal value”. Our recent study on the histopathology of OI bone demonstrated lamellar bone in both the relatively severe progressively deforming variants as well as the benign autosomal dominant variants (Shapiro et al, 2021). We also documented the more cellular nature of both woven and lamellar OI bone compared to normal human bone; this in effect reflecting the diminished matrix synthesis in OI even though the cell components are actively attempting synthesis.

### Important technical considerations in quantifying bone lamellae thicknesses using polarization microscopy

#### 1. General considerations

A major reason polarization microscopy has not been widely adopted for quantitation is the variability of findings with its use. In bone, these difficulties include: i) the wide array of angulation of the collagen fibrils of bone even beyond the obvious woven bone components where fibers in random array lead to an isotropic appearance (no light transmission). Fibrils positioned along the long axis of the cortex of bone in the lamellar conformation are not oriented with some invariably parallel to the longitudinal axis and others invariably transverse and in parallel array at right angles (in what is described as a strict orthogonal array); rather the alternate layers are best described as having unidirectional fibers rotated into twisting arrays at variable angles to one another and in densely packed and then loosely packed conformations such that the boundaries seen by polarization are not necessarily rigidly sharp. When standardized precautions are taken however, lamellation is evident and the adjacent lamellae are alternately bright and dark as shown clearly by qualitative polarization; furthermore, our study shows statistically significant measurement differences with quantitative PLM in normal and OI bone. ii) When viewing bone histology sections through a polarizing microscope one must also consider the possible effects on the polarization caused by the embedding agent (paraffin, plastic, etc.) and the chemical nature of the stain used to enhance the appearance of the bone tissues, both of which can affect the passage of light. The values and limitations of polarizing microscopy for extracellular matrix assessment have been clearly reviewed by Módis (1991).

#### 2. Specific considerations

The main criteria regarding which lamellae are measured have been outlined in Methods above. Where quantitative studies on bone are to be done, standardization of tissue preparation is highly advisable (Reid, 1986). This can include demineralization, the same embedding medium (plastic with our study), tissue sectioning at the same thickness (5μm in this study), and the same staining regimen (1% toluidine blue, a metachromatic dye) to provide a uniform tissue environment. Other tissue processing methods demonstrate polarization well; it is the standardization we stress, not necessarily, for example, the embedding or staining technique. Other reports have discussed the technical problems referable to the accuracy of lamellar measurements. Of particular importance are the thickness of the sections cut where variability can lead to altered lamellar thicknesses (Boyde et al, 1984; Kragstrup et al, 1983) and the significant effect of obliquity on lamellar thickness measurements, generally by increasing the widening (Reid, 1986).

### Contribution of findings of lamellar thickness abnormalities in OI bone to an understanding of its histopathology

#### 1. Diminished lamellar thicknesses in OI is an additional histopathologic finding for correlative assessment with collagen and related gene defects

A wide spectrum of collagen and collagen related gene mutations underlies the OI disorder (Forlino et al, 2011; Marini et al, 2017). Since these mutations affect both the structure and the amount of collagen synthesized, they are reflected in the amount and structure of the bones that form in those affected. Depending on the mutation, the OI disorder can be sufficiently severe at one end of the spectrum that the child is either stillborn or dies shortly after birth (lethal perinatal, Sillence type II, woven bone only formed) or sufficiently mild that there is full life expectancy with no deformity, the ability to walk and only occasional fractures (benign autosomal dominant, Sillence type I, lamellar bone often of normal appearance). The patients assessed in this study were sufficiently involved to have several fractures in childhood that invariably required extensive orthopedic management eventually involving surgical intervention for correction of bone deformity and minimization of fracture by osteotomy and intramedullary rod insertion (progressively deforming Sillence type III, generally with diminished lamellar bone synthesized on some persisting woven cores). In this group of patients, the lamellar bone formation was imperfect. Knowledge of the histopathology of lamellar bone in OI is crucial to an ability to appropriately quantify the lamellae by PLM. The flattened elliptical shape of mature osteocytes in normal lamellar bone has been well defined, as has the fact that osteocytes in OI bone tend to a more oval shape in both animal (oim mouse model) (Carriero et al, 2014) and human (Shapiro et al, 2021) forms. The orientation of the canaliculi also clearly aids in defining the orientation of the lamellae, since canaliculi passing at right angles from the side walls of the oval/elliptical osteocyte and at right angles to the lamellae are good indicators that the lamellae have been sectioned without obliquity along their longitudinal or transverse axes (Shapiro, 1988; Kerschnitzki et al, 2011; Shapiro and Wu, 2019). In considering the hierarchy of bone structure, it is evident that OI fragility is a cumulative problem beginning with collagen mutations and affecting structural integrity at each successive hierarchical level. OI bone has been found to be hyper-mineralized, relatively hypercellular (due to decreased matrix synthesis), and showing at the histologic level failure of normal progression along the ‘woven bone synthesis by mesenchymal osteoblasts to lamellar bone synthesis by surface osteoblasts’ pathway (Vanleene et al, 2012; Roschger et al, 2008; Shapiro et al, 2021). Correlation of collagen mutations with these findings as well as lamellar thicknesses should add further prognostic value to inform patient management.

#### 2. Value of manual and automated thickness averaging method for lamellar assessments in normal and pathologic bone

The distinct findings of diminished lamellar thicknesses in OI compared with normal bone indicate a role for this histomorphometric measurement in association with PLM in assessments of OI bone. These can be done either in diagnostic workups using transiliac biopsy or, as indicated by this study, in cortical bone fragments removed from long bones as part of corrective surgical procedures that are commonly performed. Most studies of lamellae in human bone reported above used techniques too complicated for routine pathology testing. The use of decalcified bone processed for light microscopy, however, is a straightforward clinical laboratory procedure. The widely used standardized outline for clinical histomorphometry, composed of several dozen measurements, lists lamellae as a normal component of bone tissue (their table 1) but makes no further reference to its use as a quantitative tool in its table 2 (sources and referents), table 3 (primary measurements), table 4 (derived indices) or table 5 (summation values) (Dempster et al, 2013). We suggest that lamellar thickness measurements could be included beneficially in assessments of OI bone (and other metabolic bone disorders).

### PLM Quantitation of lamellae in bone and its relation to the study of the hierarchy and mechanobiology of bone

#### 1. Studies on the structural hierarchy of bone

Bone is widely recognized as a hierarchical composite material that begins with collagen synthesis as 3 chains of amino acids twist into a triple helix. Descriptions of this hierarchy have been progressively widened and made more specific over the years (Lakes, 1993; Weiner and Wagner, 1998; Rho et al, 1998; Fratzl and Weinkamer, 2007; Reznikov et al, 2014b; Reznikov et al, 2018; Buss et al, 2022). Buss et al (2022) define 12 levels of structure, beginning with collagen synthesis and finishing with the entire skeleton. PLM, while clearly defining bright and dark layers of lamellae, cannot by itself further define the internal structure of such layers; PLM has spatial resolution at 250 nm while SEM resolution is at 1nm and TEM resolution 0.1 nm (Georgiadis et al, 2016). Many imaging techniques, currently applied to assess bone ultrastructure, have been outlined in detail by Georgiadis et al (2016). Boyde and Hobdell (1968) first showed the value of SEM in revealing the structure of lamellar bone. Shah et al (2019) have recently defined the contribution of SEM over the past 50 years in determining bone structure. While study at many of the hierarchical levels of normal and OI bone is advancing well, assessment of lamellar thicknesses by quantitative PLM can add to correlations with other levels of findings.

#### 2. Improved structural definition of bone and its relation to mechanical outcomes

Lamellar bright and dark layer thicknesses as well as their orientation and composition would be expected to play a key role in the micromechanical aspects of bone structure. This work documents how quantitative PLM provides new information on the underlying bone histopathology in OI. Beginning several decades ago, the biomechanical studies of Ascenzi and colleagues on isolated single osteons of normal human bone allowed them to assess their tensile, compressive, shearing, torsional and bending properties; these findings have been summarized and extended with updated techniques and are used to correlate with various imaging and genetic features of bone (Ascenzi et al, 2003; Ascenzi and Roe, 2012). Casari et al (2021) have assessed microtensile properties and failure mechanisms of cortical bone at the lamellar level. Micromechanical studies also involve nanoindentation at osteonal and occasionally lamellar levels (Rho et al, 1999; Carnelli et al, 2013); multiscale computational modeling of bone (and cartilage) mechanics and mechanobiology (Bolger et al, 2019; Wang et al, 2019); and theoretical and molecular modeling at the collagen fibril level (Buehler, 2006; Delpalle et al, 2015).

## Competing Interests

All authors declare no conflicts of interest.

## Funding Sources

Not applicable

## Summary

Lamellar bone that forms in moderate and severe OI is shown in this study for the first time to be composed of thinner and less regularly structured lamellae than those in normal bone. PLM histomorphometry shows mean lamellar thicknesses (bright and dark merged) are statistically significantly decreased in OI compared to normal bone as are bright and dark lamellar thicknesses measured independently. We report on the development of an automated thickness averaging method for measuring lamellar thicknesses that is, most likely, more accurate since it averages a greatly increased number of measurements per individual lamella. Lamellar thickness measurements can be helpful in assessing the effect of specific collagen and collagen related gene mutations on OI bone synthesis and warrant inclusion in research and clinical histomorphometry.

## Notes

### Competing Interest Statement

The authors have declared no competing interest.

